# A stochastic model of homeostasis: the roles of noise and nuclear positioning in deciding cell fate

**DOI:** 10.1101/2021.01.03.425080

**Authors:** Amit Jangid, Suriya Selvarajan, Ram Ramaswamy

**Affiliations:** School of Computational and Integrative Sciences, Jawaharlal Nehru University, New Delhi-110067, India; Tata Institute of Fundamental Research, Mumbai 400005, INDIA; Department of Chemistry, Indian Institute of Technology, New Delhi-110016, India

**Author notes:** These authors contributed equally to this work (Amit Jangid, Suriya Selvarajan). Electronic address (Amit Jangid).

**Keywords:** Stem cell, nucleus position, noise, cell fate, stochastic process, gene regulatory network (GRN), symmetric and asymmetric cell division, and clonal competition

## Abstract

We study a population based cellular model starting from a single stem cell which stochastically divides to give rise to either daughter stem cells or differentiated daughter cells. There are three main components in the model: nucleus position, gene-regulatory network, and stochastic segregation of transcription factors in daughter cells. We study the proportion of self-renewal and differentiated cell lines as a function of the nucleus position, which in turn decides the plane of cleavage. Both nuclear position and noise play an important role in determining the stem cell genealogies and these results can be compared with a Markov model that ignores nucleus position. We have observed long and short genealogies from model simulation which compares well with experimental results from neuroblast and B-cell division. Symmetric cellular divisions were observed when the nucleus is apical, while asymmetric division occurs when the nucleus is towards the base. The number of clones decreases as a function of time in this model, although the average clone size increases.

## Introduction

One of the fundamental question of developmental biology is to uncover the secret of how genetically similar cells give rise to distinct fate during development to precise balance between self-renewal and differentiation which in turn maintain homeostasis. One prominent mechanism for maintaining long-term homeostasis is through asymmetric cell division that yields one stem and one differentiated daughter cell at each division [1, 2]. The alternative is either uncontrolled growth with possible tumorigenesis, or tissue degeneration [3–5]. Symmetric cell division, on the other hand, leads to two daughter cells that have the same fate, namely both stem or both differentiated [6–8]. Two distinct regulatory pathways appear to play a role in deciding whether daughter cells are self-renewing or differentiated [9–11]. Specific proteins in the intrinsic mechanism determine cell fate by segregating unevenly in daughter cells that evolve stochastically [2, 12, 13]. Alternately, external factors can decide cell fate by delivering short-range molecules (proteins) [14] that communicate between cells and that can activate or block specific transcriptional networks that decide cell fate [2, 14, 15]. Through the so-called intrinsic mechanism [16], daughter cells appear to be different in shape, size, and concentration of transcription factors while in the niche mechanism [14, 15] daughter cells have the same shape, size, and transcription factors. Cell division is an extremely well-studied and complex process that consists of several steps [17, 18]. A polarity axis is set up due to the localization of specific types of proteins along the apical-basal axis of cell [12, 13, 19–21]. This axis controls the interactions between microtubules and the cortex, resulting in differential torques on the mitotic spindles and this leads to different shapes and orientation of the spindles [16, 21–24]. Transcription factors segregate differentially in different parts of the parent cell, and this results in different concentrations of proteins in daughter cells [21, 25– 29].

Cell-fate is controlled by a combination of gene-expression and intrinsic or extrinsic noise. The first suggestion, made by Waddington [30] was to consider the state of the cell as dynamical attractor [31–38]. The regulatory network has several attractors in the phase space, each corresponding to a different cell type, and each cell type is decided by a different set of genes/proteins at the population level. From a dynamical systems point of view, therefore, differentiation is considered as a transition from one attractor to another, driven by several factors. In a recent study in an early mouse embryo, it was observed that position of the nucleus can alter cell fate by controlling the division plane [39, 40]: when toward the apical side, the divisions tend to be more symmetric while more asymmetric divisions are observed when the nucleus moves towards the basal portion. This could be due to the different shape and size of daughter cells having a different concentration of expressed proteins [16]. In the absence of intrinsic noise, proteins are expressed in a correlated manner [41] whereas the same was not observed in the presence of intrinsic noise. Reducing the noise in ComK, protein that regulates competence for DNA uptake, decreases the number of competence cells [42]. Taken together, these observations suggest that these three factors, namely noise, nucleus position and the gene regulatory network (GRN) play a crucial role in deciding cell fate.

In the present work we propose a population-based stochastic model of cell division starting from a single stem cell. The three main components of the model are the nucleus position, the stochastic partitioning of transcription factors and gene regulatory network (GRN). We study the effect of factors that lead to stochasticity in the nucleus position and in the evolution and partitioning of transcription factors in daughter cells. Our aim is to understand the statistics of genealogies, namely the proportion of self-renewal and differentiated cell lines as a function of the nucleus position, and stem cell behaviour deriving from different planes of cleavage. We also study the average number of clones and average clone size as a function of time [43–45]. Our dynamical model results are further compared with a coarse-grained Markov model which ignores nucleus position. To keep the model simple we did not incorporate niche signalling and de-differentiation.

An earlier model of stem cell division [46] has considered a cellular network in which each cell consists different chemical components and these chemical component are catalysed via chemical diffusion with limited number of nutritions. The model has proposed new hypothesis that cell differentiation, growth enhancement, and robustness in a identical cell population can be self-organized. A second model [47] includes all possible screened GRNs of genes which generate cell diversity through cell-cell interaction. In the model [47] it has observed that stem cell that proliferate and differentiate exhibits oscillatory behaviour. A noise-driven cell differentiation model [48] that uses a gene regulatory network with two mutually inhibiting transcription factors shows an increasing number of differentiated cells as a consequence of noise (both intrinsic and extrinsic) in the division plane, and in uniform distribution of cell transcription factors. Noise also leads to a new phenotype among different communities [48]. The role of noise in deciding cell fate [41, 42] by altering gene expression levels in daughter cells which either activate or block particular pathways to switch the cell identity is also well known.

In the present model we observed short, ending after few divisions and having no stem cells, and non-terminating genealogies, having non zero stem cells as leaf node, similar to experimental observations [49–53]. The probability distribution of stem cells found to be bi-modal from our stochastic model and from coarse-grained Markov model. There is an initial increase in total number of live cells followed by a decrease pattern similar to experimental observations [54]. Introducing the noise in nucleus position reduces the proportion of stem cells whereas increases differentiated cells proportion. We also found that the number of clones decreases with time, although the average clone size increases, consistent with experimental observations for different stem cells [43–45, 55, 56]. In the last we found that increasing nucleus position towards apical, leads to more daughter cells having similar fates as observed in experimental studies [39, 40].

The paper is organized as follows. In the next Section we discuss the experimental evidence for the different aspects of the model, namely the gene regulatory network as well as the internal dynamical processes such as cell growth, division, and segregation of transcription factors. In Section III, we compare the bimodal distribution of stem cells obtained from a Markov chain analysis and the present model simulation. This is followed by a summary and discussion of our results.

## Materials and methods

### Experimental evidences supporting model components

The purposed stochastic model consists of three main components, nucleus position, stochastic segregation of transcription factors, and gene regulatory network (GRN). In an early mouse embryo, more asymmetric divisions have been observed as nucleus moves towards apical whereas, more symmetric divisions were observed for apical positioning of nucleus [39, 40]. This provides an insight that nucleus position may alter the cell fate indirectly by deciding different shape and size of daughter cells with different level of transcription factors. It has observed [39] that transcription factor Cdx2 (a protein expressed in intestinal epithelial cells) regulate nucleus movement towards apical or basal by increasing or decreasing its level. Down regulation of aPKC (protein involved in cell polarity) molecule also doubles the asymmetric division with a decreasing number of apical nuclei [39]. It has suggested and also observed experimentally that cell state can be considered as an attractor on an energy lands scape and can be expressed in a mathematical form which consists of different types of transcription factors [31–38]. Such as Cdx2 *>* Oct4 induces trophoectoderm (TE) fate, whereas Oct4 *>* Cdx2 activates the inner cell mas (ICM) program [38]. Similarly, in hematopoiesis two different transcription factors, GATA1 and PU.1, regulate cell fate decision of different cell types [38]. Recently [32], high-dimensional stable attractor has been observed in which an attractor represents distinct cell fate and differentiation can be seen as a transition from stem cell attractor to different attractor by several intrinsic and extrinsic factors [32]. Collectively these studies suggest that the different transcription factors can be modeled mathematically where, different concentration of transcription factors, expressing different gene level, can guaranty of different cell identity. It has observed [41] that in the presence of intrinsic and extrinsic noise (measured by cyan fluorescent protein and yellow fluorescent protein), in a single cell, results in the variation of gene expression and in another experiment [42] it has showed that variation in gene expression can lead to transitions between alternative states of gene expression.

### Dynamical process of the model proceeds as follows

We consider the regulatory network in FIG. 1A, with a pair of transcription factors, labelled *X* and *Y* that mutually inhibit each other while being self-activating. The stoichiometric equations are straightforward to obtain (Table I) and through standard analysis it can be seen that in this system there are a large number of dynamical attractors for any set of parameters. Three are of special importance here (FIG. 1B), and they are all fixed point attractors: when the asymptotic values *X*^*∗*^ and *Y* ^*∗*^ are identical, labelled S, and when the asymptotic values of *X* and *Y* are unequal. Type A has *X*^*∗*^ *> Y* ^*∗*^, while Type B has *Y* ^*∗*^ *> X*^*∗*^. In our interpretation, Type S corresponds to stem cell, while Types A and B are differentiated cells. The basins of attraction for these three types of cells are shown in FIG. 1C for a typical choice of parameters. Other regulatory networks can also be incorporated in such models [35–37]; combinations of the toggle switch [35] and repressilator [35] which can have different steady states. Such systems show richer dynamics.

**FIG 1:**
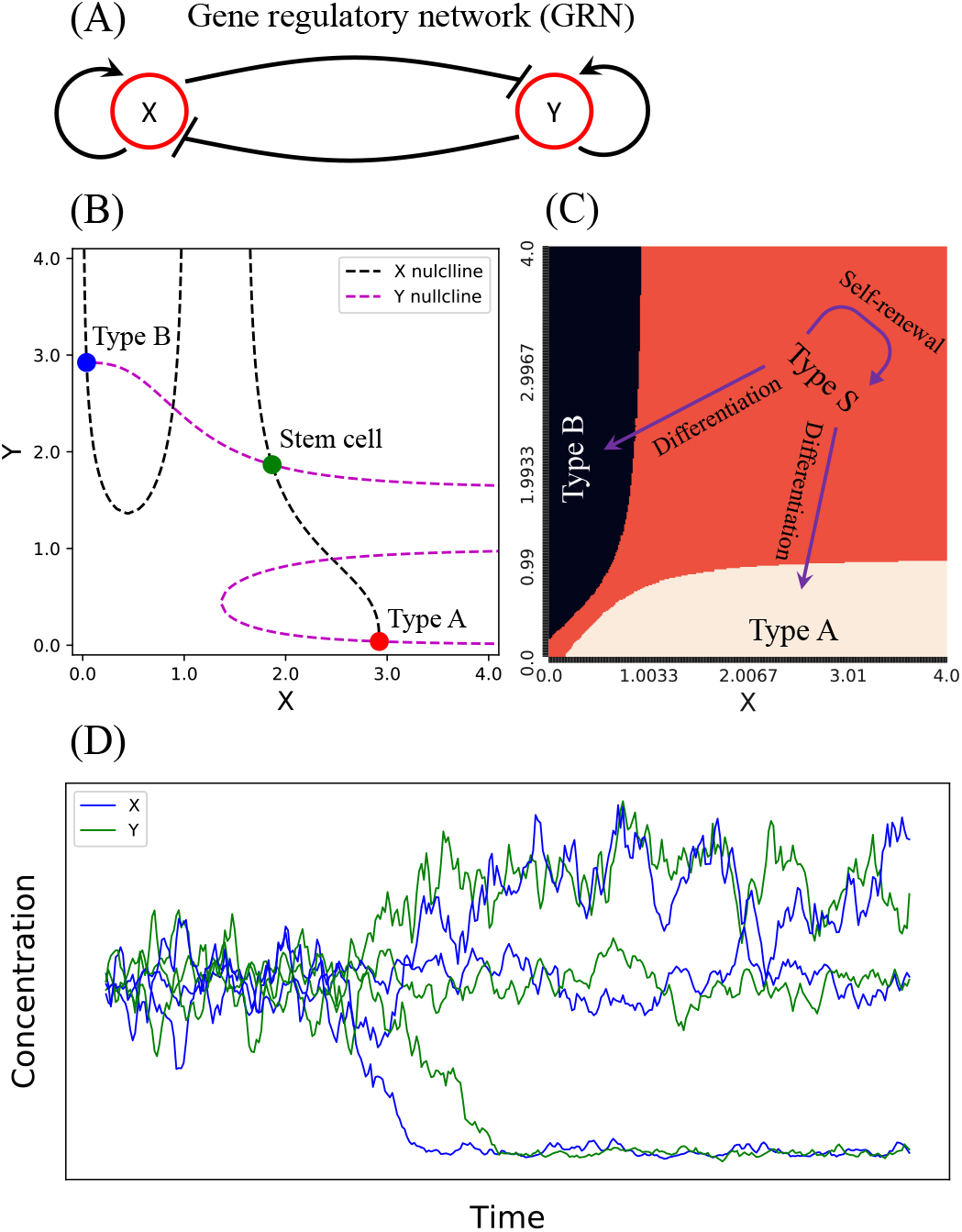
Gene regulatory network and basin of attraction: (A) Gene regulatory network with two transcription factors X, and Y inhibiting each other and self-activating. (B) Nullcline showing three different attractor corresponding to three different cell type S, A, and B. (C) Basin of attraction in which three different shaded region correspond to three different cell types S, A, and B in phase space (*X* and *Y* are concentration). Blue arrow from S to S represents self renewal, while arrow toward A, and B region correspond to differentiation due to internal or external noise. S represents stem cell, A and B represent differentiated cells. (D) shows three different time course of transcription factors X, and Y corresponding to (C).

**TABLE I:** Reactions pertaining to the regulatory network shown in FIG. 1A. *X* and *Y* are transcription factors, *S*, and *P* are substrate and products respectively.

| No. | Reaction | Rate constant |
| --- | --- | --- |
| 1. | $S \rightarrow X$ | $r_1 = \alpha_1 \frac{X^n}{1+X^n} + \beta_1 \frac{1}{1+Y^m}$ |
| 2. | $S \rightarrow Y$ | $r_2 = \alpha_2 \frac{Y^m}{1+Y^m} + \beta_2 \frac{1}{1+X^n}$ |
| 3. | $X \rightarrow P$ | $r_3 = \mu_X$ |
| 4. | $Y \rightarrow P$ | $r_4 = \mu_Y$ |

The dynamical equations corresponding to the above model are studied in the usual manner, starting from an initial volume *V*_0_, a single initial stem cell with nucleus at position *P*_0_, and a random initial concentrations of *X* and *Y* (corresponding to stem cell attractor). Assuming the presence of sufficient chemicals for proper cell growth, the cell can be taken to grow at an exponential rate while in the absence of adequate nutrients, the rate of growth seems to be linear. We have used Gillespie algorithm [57] in our simulations where volume depends on time (*V* =*V*_0_*exp*^*λt*^). The decay rate *µ* is independent of volume, but the other rates (autocrine and paracrine terms) are volume dependent (see Table). The volume dependent rates give rise to additional complexity in the dynamics [57].

At the time of cell division, the plane of cell cleavage is decided by the position of the nucleus and the segregation of transcription factors [17, 18, 39, 40]. The parameter *P* (see Fig. 2 top box) is the fraction of the distance along the principal axis of the cell (measured from the base) at which the nucleus is situated. As soon as the volume reaches a threshold *V*_*C*_, the cell divides, with probability *P* along the vertical plane, and with probability 1-*P* along the horizontal. Transcription factors are also distributed stochastically in the two daughter cells, proportional to the relative volumes, and the nucleus position is set at random (Appendix) in the newly created daughter cells as well. Starting from a single stem cell as the root of the genealogical tree, the system is propagated in time as follows. The cell grows in size as described above and once the volume exceeds *V*_*C*_, division takes place. Eqs. 2,3 of the regulatory network are used for the evolution of transcription factors in each of the daughter cells. If the daughter cells are stem, then further division can take place. Data for each cell is collected and summarized in a genealogical tree with averaging over 500 cell divisions; see FIG. 2 for a representative result. The precise rules of cell division are specified in the Appendix.

**FIG 2:**
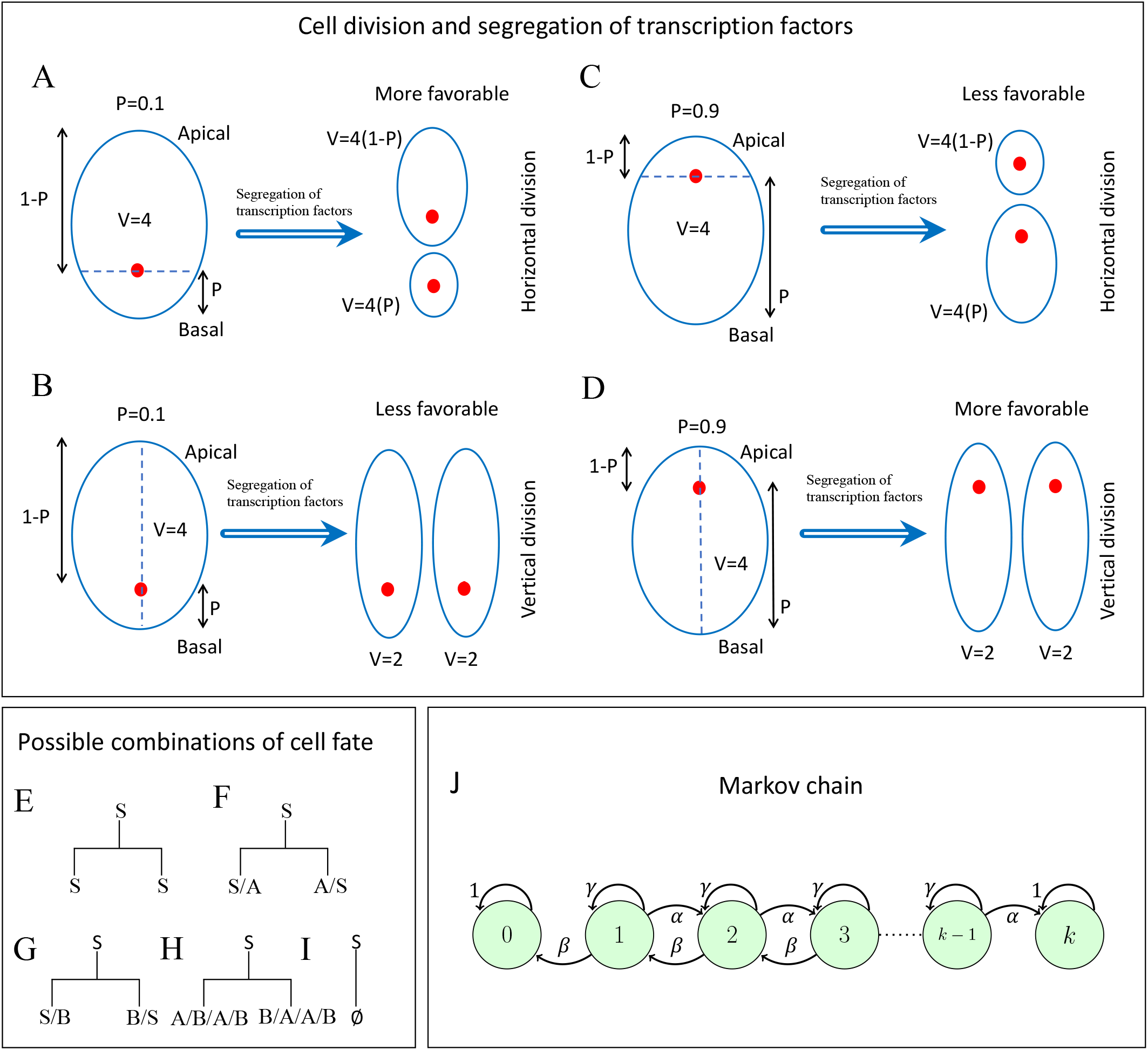
Cell division and segregation of transcription factors. (top box). Schematic diagram (top box) of a stem cell for horizontal and vertical division for different nucleus position and segregation of transcription factors in daughter cells. Horizontal and vertical dashed (blue) line through nucleus (red dot) correspond to horizontal and vertical division of cell respectively. *P* is the height of nucleus from basal, and 1− *P* is from apical. For *P* = 0.1, horizontal division (A) is more favourable than vertical division (B), whereas for *P* = 0.9, vertical division (C) is more favourable than asymmetric division (D). **Possible combinations of daughter cell fate in each cell division** (bottom left box). (E) and (H) correspond to symmetric renewal and symmetric differentiation respectively whereas (F) and (G) correspond to asymmetric division and (I) corresponds to cell death (though we did not count cell death throughout the model). S indicates stem cell, A and B indicate differentiated cell, and *φ* indicates dead cell. **Markov chain** (bottom right box). The schematic diagram (bottom right box) represents Markov chain for cell division starting from single stem cell. *k* corresponds to the number of stem cells. *α* is the probability of getting both daughter cells as stem cell (+1 state), *β* is probability of getting both cell as differentiated cell (−1 state), and *γ* is the probability being in the same state (no changes) with *α* + *β* + *γ* = 1. Probability of being in the same state is 1 for state *k* = 0, and *k* (these two states are absorbing states).

The average clone size and clone number as a function of time is also monitored. We consider a simple case of clonal competition in which a stem cell is replaced, based on it’s division time, in the different pool of clones. The stem cell with the earliest time of division is stochastically lost, and will be replaced by daughter cells, via symmetric renewal (gain in stem cell by one), symmetric differentiation (loss in stem cell by one), and asymmetric renewal (no loss or gain). In this competition, there is no interaction among members of the set.

### Markov chain

We can device a Markov model [58, 59] for cell division as shown in FIG. 2J, where *α* is the transition probability from state *q*_*i*_ to state *q*_*i*+1_ when both daughter cells are stem. Correspondingly, *β* is the transition probability from state *q*_*i*_ to state *q*_*i−*1_ when the daughter cells are differentiated cells, and *γ* is the stay put transition probability which arises when one daughter cell is stem cell and the other is differentiated. We have a set of states, *Q* = {*q*_0_, *q*_1_, *q*_2_ … *q*_*k*_}, where *q*_*i*_(*i* ∈ (0, *k*)) being a state having *i* number of stem cells and also *Q*_0_ = {0, 1, 0 … 0} as we always start from a single stem cell. State *q*_0_ and *q*_*k*_ are absorbing states such that it is impossible to leave these states (*α, β, γ* = 0.0, 0.0, 1.0) and the transition probabilities from state *i* to *j* is 0∀ *j* ≠ *i*± 1. So the transition probability matrix for the Markov model is as follows,

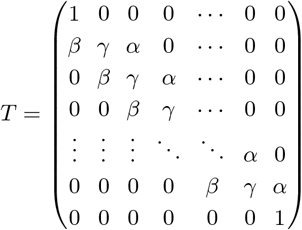

where raw and column represents states from *q*_0_ to *q*_*k*_. Probability being in state *q*_*j*_ from state *q*_*i*_ after n divisions is *P*_*i*+1,*j*+1_ element of *T* ^*n*^ ∀ *i, j* ∈ (0, *k*). In this model *i* is always fixed at 1 as we have started from single stem cell, so each element of second raw in the matrix *T* ^*n*^ corresponds to the probability of getting state *q*_*j*_ (*j* = 0 … *k*) from *q*_1_ (having one stem cell).

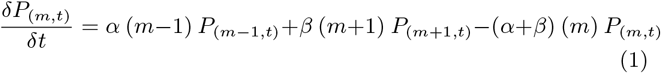

Probability evolution of having *m* stem cells at any instant of time *t* can also be calculated from master equations as shown in Equ. 1. Here, 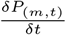 represents the rate of change in the probability of having *m* stem cells at time *t*. First term in right side refers to the overall probability coming from state *m* −1 to state *m*, second term from *m* + 1 to state *m*, and last term refers to leaving the state *m* either to state *m* + 1, or to state *m* 1. −*P*_(*m,t*)_ is the probability of having *m* stem cells at time *t*.

## Results

We study the dynamics of the proposed population model for different nucleus positions such as 0.1, 0.5, and 0.8. We have taken four different cases of cell division namely *horizontal, vertical, mixed*, and *random*. 1) *Horizontal* : in each cell division, plane of cleavage will be horizontal through nucleus. 2) *Vertical* : irrespective of the nucleus position plane of cleavage will be vertical. Each daughter cell will be having half of mother cell volume. 3) *Mixed case*: in each cell division plane of cleavage will be decided by nucleus position *P*. Volume and transcription factors will also be decided by nucleus position. 4) *Random case*:

In this case, horizontal and vertical both will be having same probability. We have also introduced noise in nucleus position in each case and compared the results.

## Markov simulation

### Bi-modal distribution of stem cell

We have studied the probability distribution, and the population of stem cells as a function of division count for different set of parameter values *α, β*, and *γ*. Probability distribution, FIG. 3A,B,C, which shows a bi-modal distribution of stem cell for different parameter set (*α, β, γ* = 0.4, 0.2, 0.4), (*α, β, γ* = 0.6, 0.15, 0.25), and (*α, β, γ* = 0.8, 0.1, 0.1) respectively. Increasing *α*, and decreasing *β* results in the increase of stem cell population whereas decreasing *α*, and increasing *β* give lower number of stem cells. It might be due to increase in the number of short terminated trees with no more stem cells available to further divide. Increasing *γ* corresponds to no changes in stem cell population but observed more number of differentiated cells because of asymmetric cell division as it provides one differentiated cell with one stem cells. In short, parameter *α, β*, and *γ* represent symmetric renewal, symmetric differentiation, and asymmetric division respectively. Stem cell population as a function of division count has shown in FIG. 3D, E, and F for same parameter sets respectively. Red curve represents stem cells population which terminate after few cell divisions and blue curve represents population which grow and does not terminate. There are 17, 9, and 3 terminated red curve and 83, 91, and 97 non-terminated blue curves in FIG. 3D, E, and F respectively (taken 100 random samples). The stem cell distribution shown in FIG. 3A, B, and C are taken from 5000 samples. Each sample is generated by evolving random number for different set of parameter values. A similar probability distribution can be obtained from transition probability matrix and master equation (Markov chain). The short and non-terminating genealogies observed in experiments [49–51] suggest that it may also provide similar bi-modal behavior. In which short genealogies will correspond to the first mode having 0 stem cell (all leaves nodes are differentiated cells), and non-terminating genealogies correspond to the second mode having non-zero stem cells.

**FIG 3:**
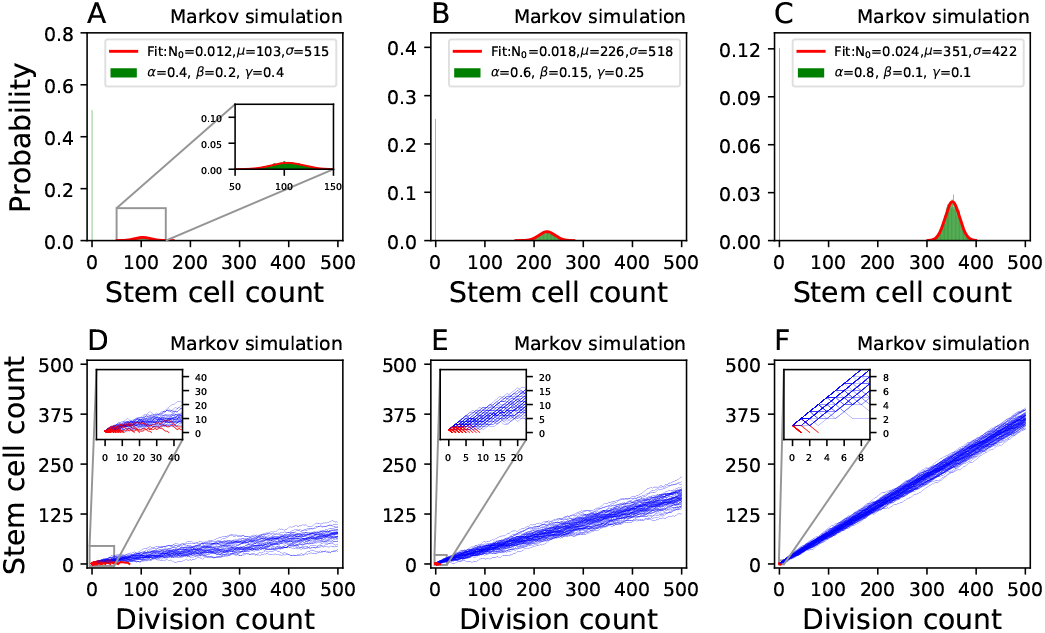
Probability distribution of stem cells obtained from Markov process. Top row, A, B, C, refer the probability distribution of stem cell for three different set of *α, β*, and *γ* averaged over 5000 samples. The red curved is the Gaussian fit for standard parameters N_0_, *µ, σ*. Bottom row, D, E, F, represent stem cell population as a function of cell division count (taken 100 random samples) corresponding to the parameter used in top row. Red color represents terminated cell lineages, and blue shows nonterminated lineages. Each cell lineage has stared from single stem cell. Inner panels in A show the zoomed in image of particular region.

## Model simulation

### Short and non-terminating genealogies

First short and long non-terminated genealogies were observed in central nerves system with symmetric and asymmetric tree [49, 50]. Long (non-terminated) genealogies were the results of symmetric division (self-renewal) whereas short genealogies were the results of asymmetric division or symmetric differentiation. The purposed model also mimics similar behavior of genealogies for different nucleus position, plane of cleavage, and noise showing symmetric and asymmetric division pattern (FIG. 4). Red, blue, and green nodes correspond to type A, type B, and stem cell respectively. FIG. 4A-C are the genealogies terminating just after first generation in which FIG. 4A, and B are symmetric having differentiated daughter cells with similar fate whereas FIG. 4C is asymmetric with two different differentiated daughter cells. FIG. 4D-I are the genealogies terminating after two generations being FIG. 4D-F asymmetric, and FIG. 4G-I symmetric. FIG. 4J-M are terminating trees after three to five generation being all asymmetric. While rest, FIG. 4N-T, represent non-terminating genealogies having stem cells as leaves nodes. As tree sizes increase it is very less probable having symmetric genealogies due to three different cell types. Few genealogies go beyond 50 generations (in case of horizontal division with noise in nucleus position). We have also observed that vertical division leads to more non-terminating genealogies than horizontal division because of more stem cells. In every short terminated genealogy there is always 0 stem cell.

**FIG 4:**
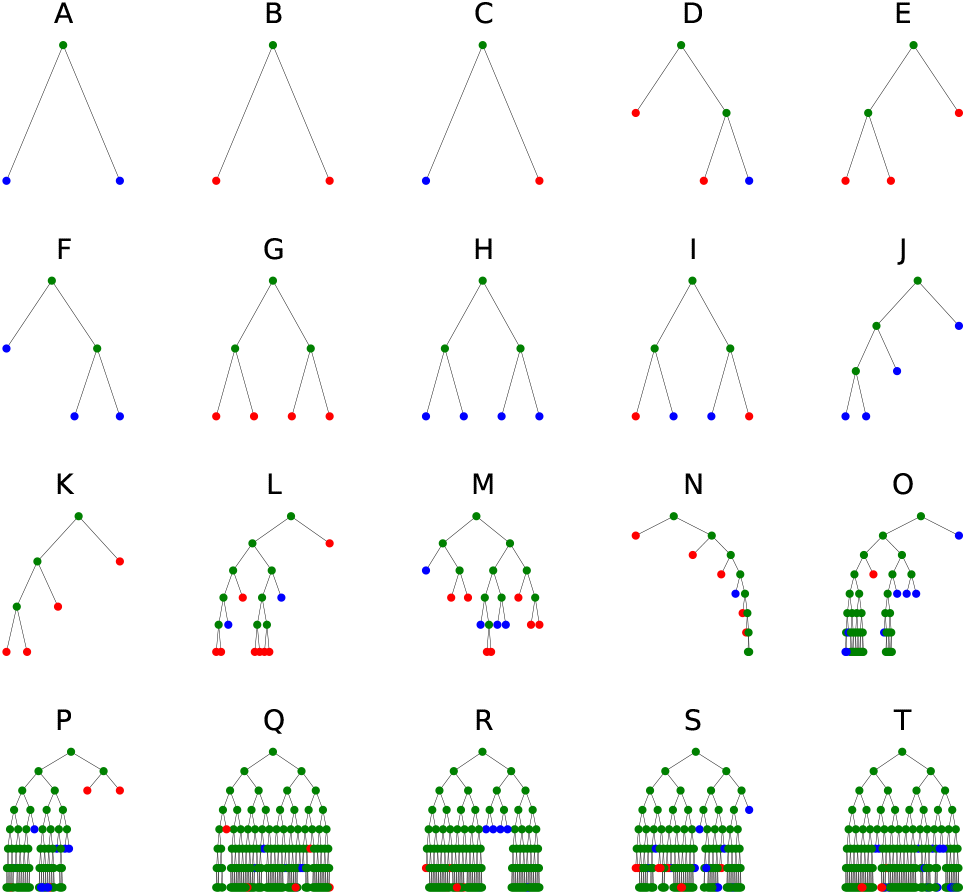
Genealogies obtained from model simulation. Short terminated and non-terminated genealogies pattern for different cases with symmetric and asymmetric division. Panel A-M are the short terminated genealogies, while rest are the larger genealogies which further divide. We have shown here maximum 7 generations. Red, blue, and green nodes correspond to cell type A, and type B, and stem cell respectively.

### Bi-modal distribution of stem cell

Observing short and non-terminating genealogies from experiments suggests that there is possibility of two modes of cell distribution each with different mean and variance. We have plotted stem cell distribution for different cases which shows bi-modal distribution similar as observed from Markov process (FIG. 5A-C), and also observed similar pattern of stem cell population as a function of division count (FIG. 5D-F). From coarse graining we have calculated three sets of *α, β, γ* such as (0.416, 0.279, 0.305), (0.557, 0.223, 0.22), and (0.813, 0.088, 0.099) from 5000 samples (FIG. 5A, B, and C respectively). For same three sets of *α, β, γ*, we have plotted probability distribution from Markov process (FIG. 5G, H, and I respectively). Stem cell distribution in FIG. 5A, and B are in very good agreement with distribution in FIG. 5G, and H in the probability of stem cell, and in the range of cells numbers (fitted Gaussian distribution), whereas distribution in FIG. 5C gives good agreement in the cell numbers with panel FIG. 5I but not in the probability. Cell counts, from model (FIG. 5D, E, and F), is also in good agreement with Markov process (FIG. 5J, K, and L respectively). Markov approach gives a very simple picture of cell distribution having no nucleus position, while the purposed model gives much rich behavior because of having different types of plane of cleavage, nucleus position, and gene regulatory network.

**FIG 5:**
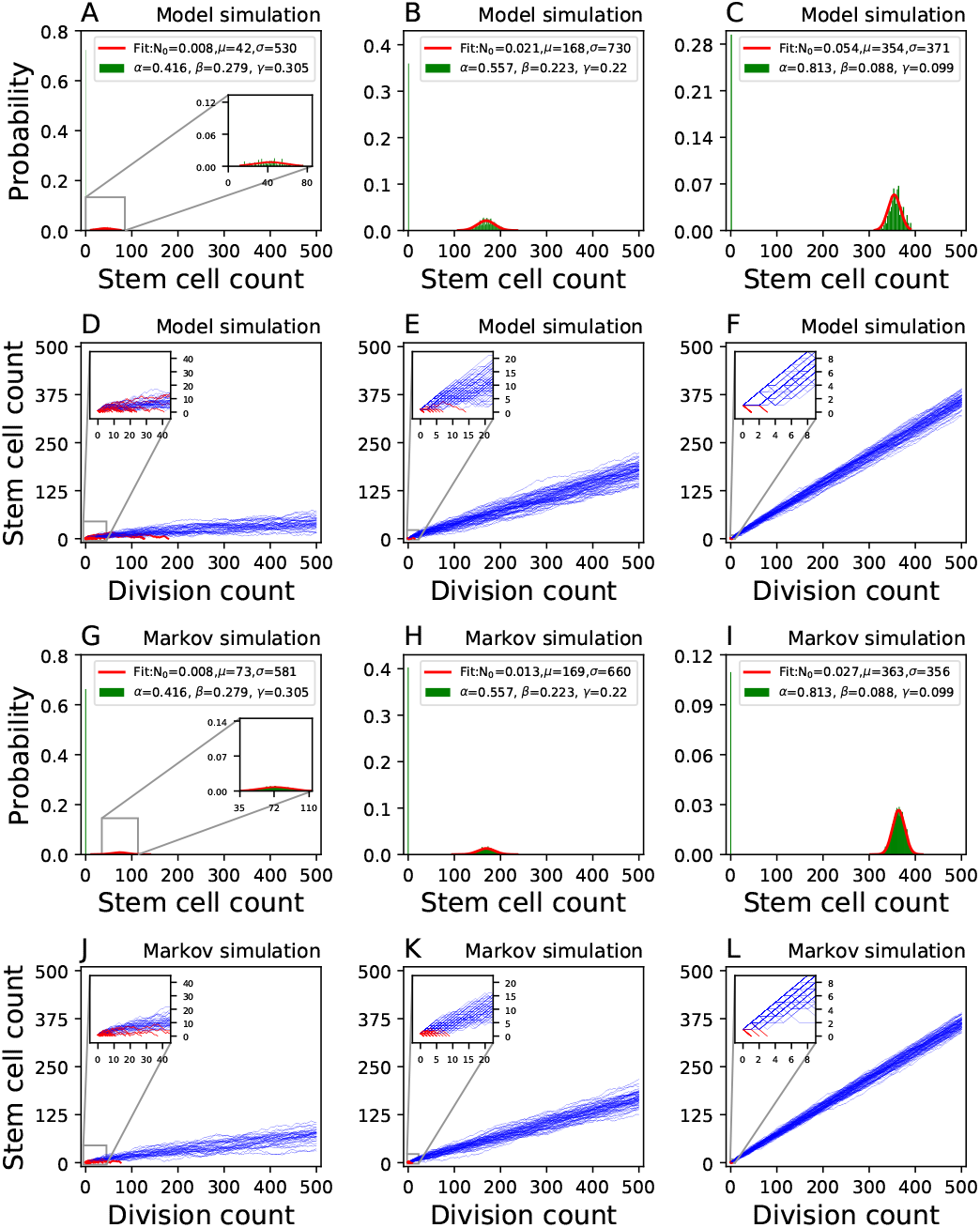
Bi-modal behavior of probability distribution from model and Markov simulation. Top row shows probability distribution of stem cell for different set of *α, β*, and *γ* calculated over 5000 samples from model simulation. Second row represents stem cell population as a function of cell division count corresponding to the parameter used in top row for 100 trials. Third row represents same probability distribution for same set of *α, β*, and *γ* from Markov simulation calculated in top row, and bottom row represents stem cell population as a function of cell division count from Markov simulation corresponding to the parameters calculated in top row. Blue and red color correspond to non-terminating, and terminating genealogies. Gaussian fit for top and third row is shown in red color for standard parameters N_0_, *µ, σ*

### Nucleus position and noise altering cell fate decision

#### 1 Cell distribution comparison among similar plane of cleavage

The proposed stochastic model and Markov process both result bi-modal distribution of stem cells which are in good agreement. We have plotted the distribution of differentiated cell type A, B, and stem cells for 4 cases (with and without noise in nucleus position) for each 0.1, 0.5, and 0.8 nucleus position (FIG. 6 only for nucleus position 0.1, see FIG. S1 and S2 for 0.5 and 0.8). In each case of cell division probability and range of cell type A against B is similar in case of noise in nucleus position such as FIG. 6A_1_ has same range and probability distribution as A_2_, same for B_1_ v/s B_2_, C_1_ v/s C_2_, D_1_ v/s D_2_. It is due to the same parameters (*n*=*m*=3.0) taken in gene regulatory network, which gives the symmetrical nature of cell type A and B from the stem cell attractor (FIG. 1B,C). In this case, the same noise level results in the same chance of transition, whereas for a different set of parameter values (*n ≠ m*), the chance of transitions biases towards one cell type than the other. Noise in nucleus position results in different stem cell range with different probabilities (FIG. 6A_3_,B_3_,C_3_,D_3_) then cell type A and B. Second peak in the cell distribution of stem cells range from 20-70 (FIG. 6A_3_), 320-400 (B_3_), 125-290 (C_3_), and 130-220 (D_3_) for asymmetric, vertical, mixed, and random case with noise in nucleus position respectively while there is sharp peak at zero (first peak) with different probabilities.

**FIG 6:**
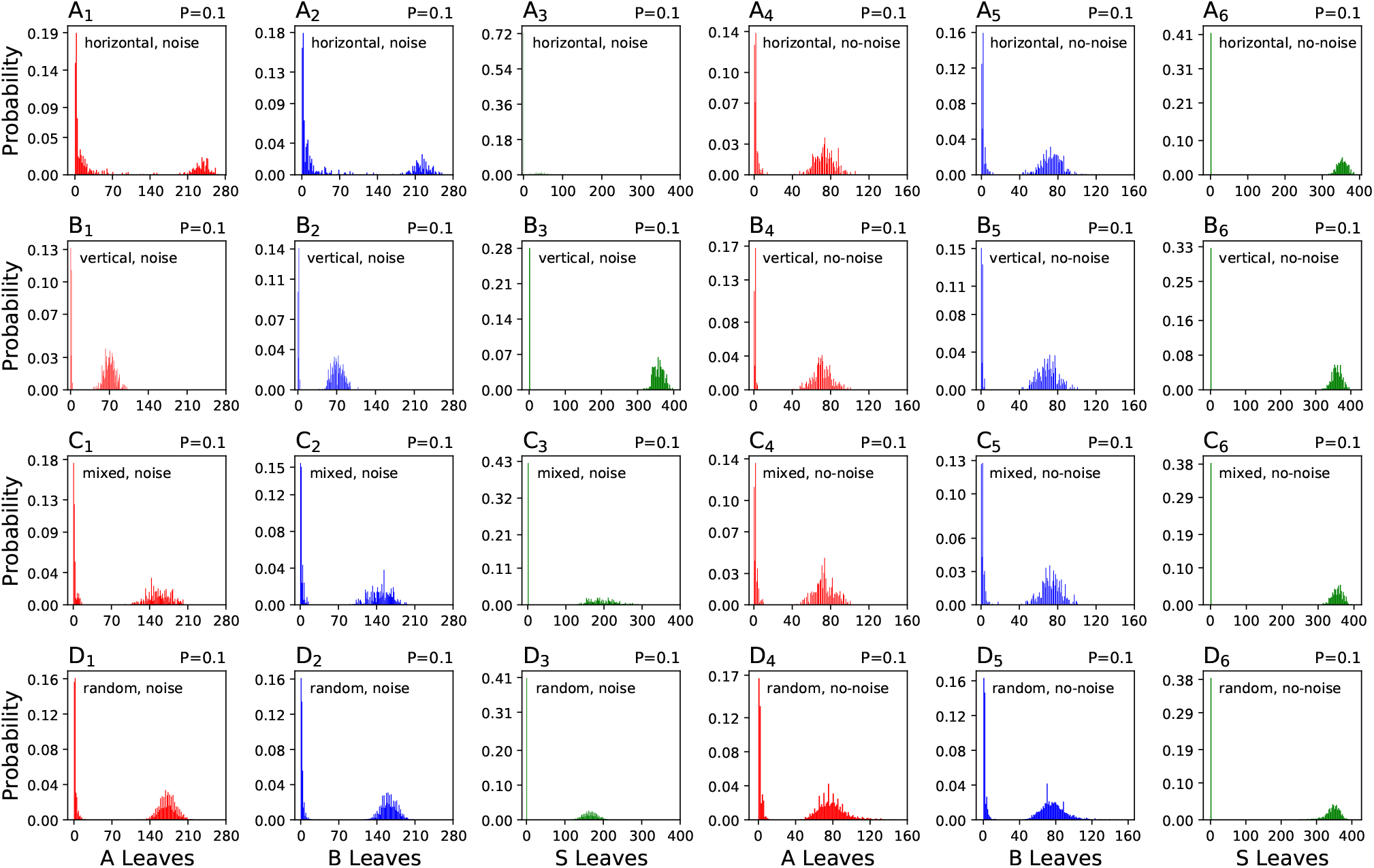
Probability distribution of differentiated cell type A, B, and stem cell S. Probability distribution of cell A (differentiated, red color), B (differentiated, blue color), and S (stem cell as leaf, green color) for all 8 different cases asymmetry, symmetry, mixed, and random with noise and without noise in nucleus position, averaged over 5000 samples. In each panel on the top right *P* represents nucleus position, and in the panel, first, and second text represent type of plane of cleavage, and noise in the nucleus position, respectively.

No-noise in nucleus position, results in the distribution with same probability and range (40-120) of cell type A against B such as FIG. 6A_4_, B_4_, C_4_, D_4_ against FIG. 6A_5_, B_5_, C_5_, D_5_ respectively. And stem cells also range (second mode) from 320-400 (FIG. 6A_6_, B_6_, C_6_, D_6_) in all four cases and also give a sharp peak at zero (first mode). These results suggest that noise in the nucleus position gives lower number of stem cells due to short genealogies, where no-noise results in long non-terminated genealogies with less differentiated cells. There is a larger range of stem cells (second mode, FIG. 6B_3_) because of noise does not change the plane of cleavage as vertical division is independent of nucleus position. Red, blue, and green color in FIG. 6 represent differentiated type A, type B, and stem cells respectively. Similar behavior has observed for nucleus position *P* =0.5, and 0.8 shown in FIG. S1 and S2, respectively.

#### 2 Cell distribution comparison among different plane of cleavage

In case of noise in the nucleus position there are different ranges of cell type A, B, and stem cells (FIG. 6A_1_-A_3_, B_1_-B_3_, C_1_-C_3_). Random case gives similar range as mixed case (FIG. 6C_1_-C_3_ and D_1_-D_3_). Horizontal division gives the largest range of cells A and B (first mode: 0-140, second mode: 140-270) with the lowest range of stem cells (first mode: 0, second mode: 20-70), whereas vertical division gives the lowest range of cells A, and B (first mode: 0-10, second mode: 40-100) with highest range of stem cells (first mode: 0, second mode: 300-400). In case of no-noise, the range of A, B, and stem cells are same across all four different cases (FIG. 6A_4_-A_6_, B_4_-B_6_, C_4_-C_6_, D_4_-D_6_).

### Fraction of stem and differentiated cell as a function of nucleus position and noise

We have studied fractions of differentiated and stem cells with different nucleus positions for eight cases (FIG. 7). We have observed the highest number of total differentiated (A+B cells) and stem cells for 0.1 and 0.5 nucleus position respectively among all nucleus position. Horizontal division with noise produces more differentiated and less stem cells (FIG. 7A). Whereas no-noise case (FIG. 7E) results in just opposite scenario with 0.5 giving maximum number of stem cells. In horizontal division, nucleus position *P* is just complementary of nucleus position 1 − *P*, hence nucleus position 0.5 leads highest stem cells than differentiated cells (due to non-terminating genealogies). In the case of vertical division with noise and without noise (FIG. 7A, F), there is no variation in number of differentiated and stem cells as equal volume and transcription factors in daughter cells. In mixed case with noise (FIG. 7C), stem cell population increase and differentiated cells decrease as nucleus position increases, but for each nucleus position, differentiated cell are higher than stem cells. Mixed case with no-noise (FIG. 7G) reverse the situation with more stem cells. In random case, with noise and no-noise (FIG. 7D, and H), there is no variability in the population of differentiated cells, and stem cell except that in case of noise, there is higher number of differentiated cells than stem cells. FIG. 7I, and J show vertical division count and symmetric fate as a function of nucleus position. Indicating that more vertical division (which is the result of increasing nucleus position) leads to more symmetric division. Here if both daughter cells are differentiated cell (either A, B or combination of A and B) or stem cells are counted in symmetric division.

**FIG 7:**
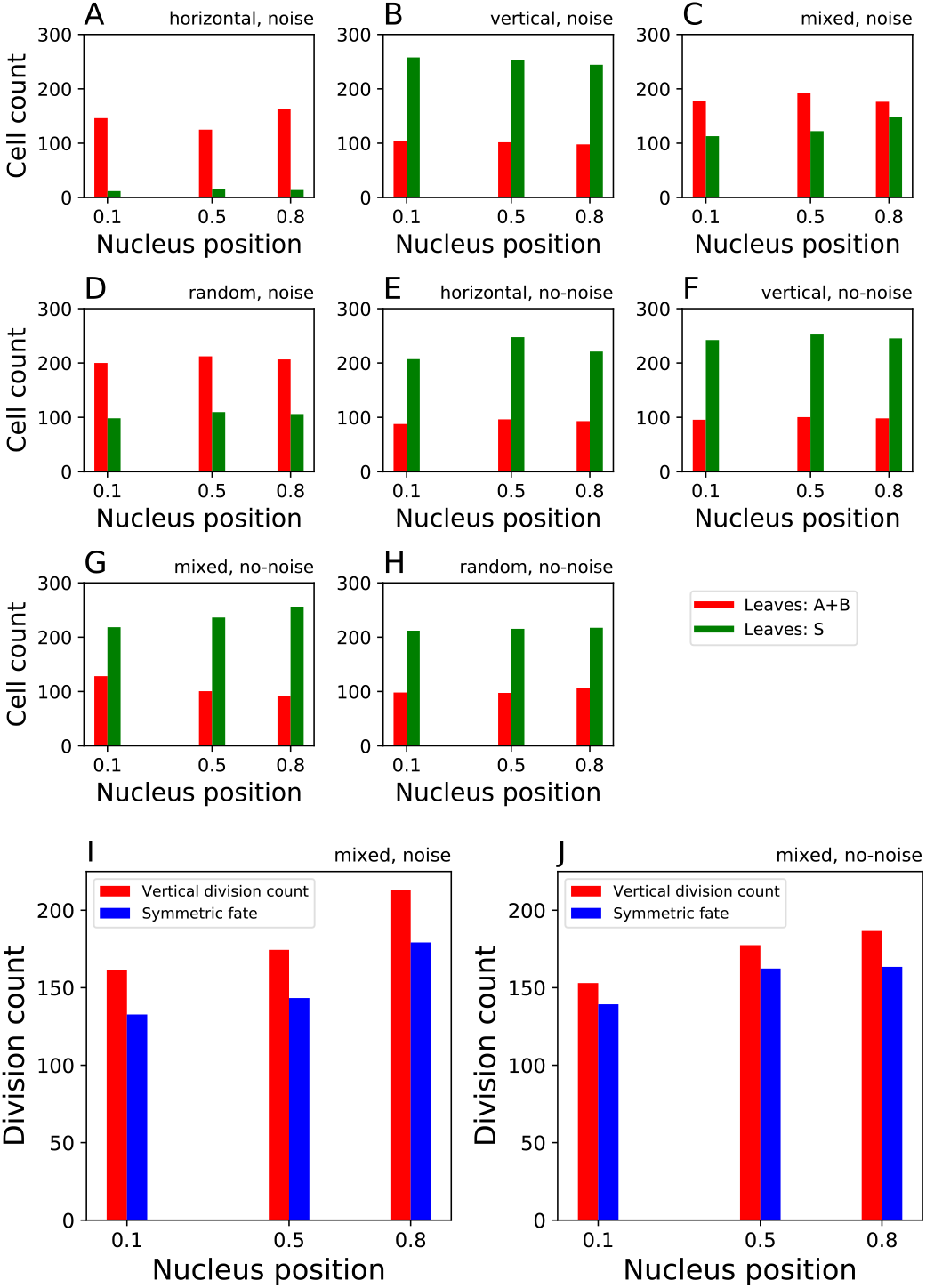
Number of differentiated and stem cells for different plane of cleavages with different nucleus position: Bar plot for number of total differentiated cells (A+B, red colour), and total S cells (leaves, green colour) for nucleus position 0.1, 0.5 and 0.8 for 8 different cases, averaged over 5000 samples. First and second text on top right in each panel represent plane of cleavage and noise in the cell division respectively. **Increasing nucleus position increases symmetric cell division**: Number of symmetric cell division (symmetric fate) from total number of vertical division for nucleus position 0.1, 0.5, and 0.8 for mixed cases with (I) and without noise (J), averaged over 5000 samples. If both daughter cells are differentiated (either cell type A, B or combination of A and B) or stem cells, are counted as symmetric division (same fate). First and second text on top right in each panel represent plane of cleavage and noise in the cell division respectively.

### Population dynamics of cell over time and generation

We have studied the cell dynamics in different generations as a function of time. FIG 8A, B, and C represent similar qualitative cell dynamics as observed in experimental studies [51, 54, 60, 61]. The dynamic shows that cell counts, green dashed line, starts increasing after initial cell divisions and reaches up to a certain level followed by a decreasing cell line, FIG 8A, B, C for three different nucleus position 0.1, 0.5, 0.8 respectively. The observations indicate that initially there are more symmetric divisions for certain time intervals, which give more stem cells to divide (overall number of cells increases) and later more symmetric differentiation and asymmetric division, which eventually leads to more specialized cells that do not divide further. For nucleus position 0.1, 0.5, and 0.8 (FIG 8A, B, C), the peak value of cell count increase as nucleus position shifts towards apical (55, 64, 81 cell count), and the time where peak has observed shifts towards left side (193, 152, 128). These results suggest that there is an increase in the number of symmetric divisions as nucleus position shifts towards apical portion. FIG 8D, E, F represent stacked area plot for individual generation, up to 20 generations, as a function of time. We have observed that cell dynamics in individual generation, FIG 8G, H, I, follow Gaussian distribution with different maximum cell number at different time, which are qualitatively similar to experimental observation.

**FIG 8:**
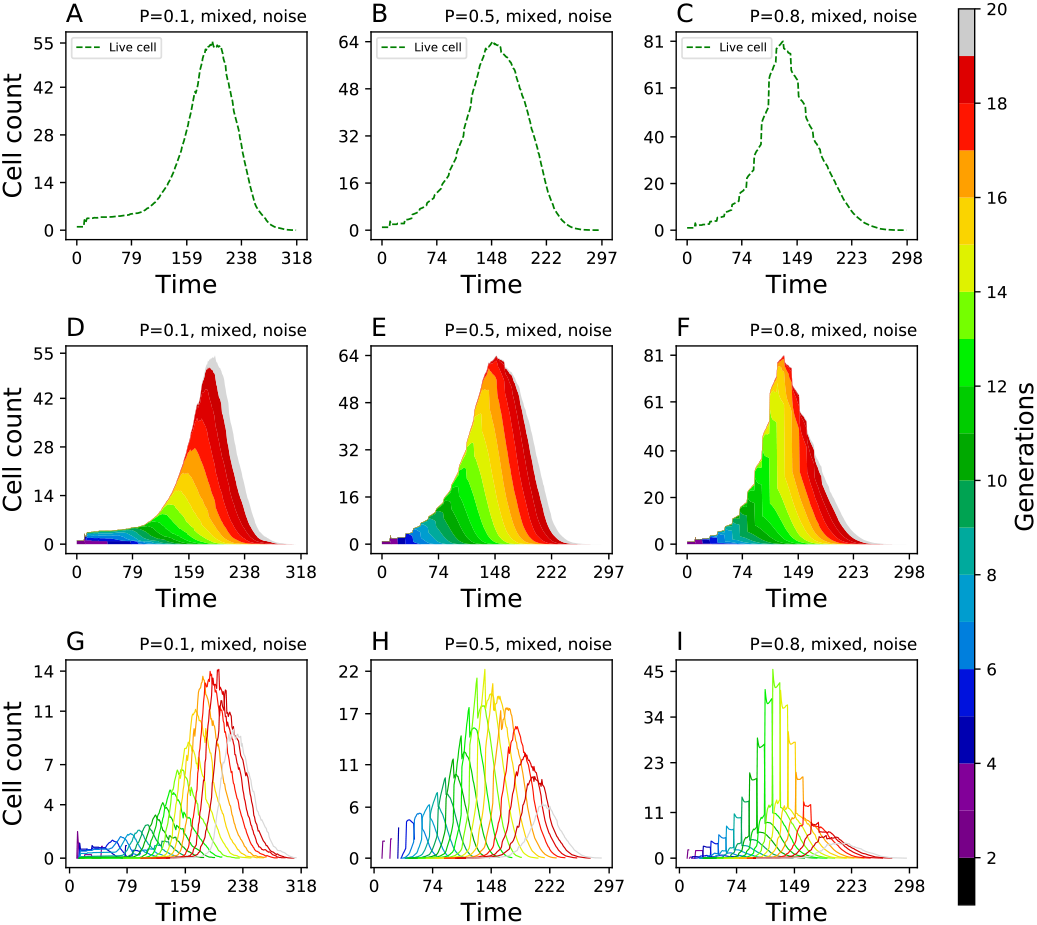
Population dynamics of cell over time. Top row displays total live cell dynamics over time. Second and third row represent stacked area plot of total cell counts with the contribution of each generation (up to 20 generations), and cell counts in individual cell generation respectively. Color spectrum represents for different generations for second and third row. In each panel on the top right *P* represents nucleus position, second, and third text represent type of plane of cleavage, and noise in the nucleus position respectively.

To study the effect of different nucleus position, plane of cleavage, and noise in the nucleus position on cell dynamics, we have plotted cell dynamics for all 24 different conditions (FIG. S3). Cell dynamics (thick black line) show common behavior across all panel, that is initially increasing pattern of cell counts followed by a decreasing pattern after attaining a certain threshold. It also suggests that the presence of the noise in nucleus position, keeping nucleus position and plane of cleavage similar, leads to more number of generations and time taken to achieve a certain population level. Such as the number of generations is going beyond 50 in FIG. S3A_1_ whereas, in FIG. S3A_5_ number of generations have observed 17 and time for these two conditions are 747 days and 235 days, respectively. Similar scenario can be observed across different panels of FIG. S3. There is no variability in the cell dynamics in each panel across second column and fourth row of FIG. S3. Vertical division is independent of nucleus position, and noise so there will be no variability in the cell dynamics (second raw), whereas in FIG. S3B_5_, B_7_ and B_8_, *P* = 0.5 behave as an attractor in which nucleus position will be 0.5 in daughter cells throughout the divisions which will lead to no variability in cell dynamics. We have also plotted cell dynamics (FIG. S4) in an individual generation for different cases, which shows Gaussian distribution. In summary, we have observed three behavior, more symmetric division for apical nuclei, symmetric division takes lesser time to achieve a certain threshold than asymmetric division, and noise increases number asymmetric division (larger number of generations are possible through asymmetric division for certain carrying capacity).

We have plotted the dynamics of maximum cell count in an individual generation for three different nucleus position (FIG. S5A, B and C), which shows a Gaussian distribution with fitted parameter values N_0_ =13.318, *µ*=16.202, *σ*=4.04, suggesting that increasing *P* (towards apical), increases the peak value of the maximum cell count however decreases the generation for that peak value. FIG. S5D, E and F represents the probability distribution of cell division showing decaying pattern (FIG. S5E showing positive skewed distribution), and FIG. S5G, H, I show fraction of undivided cells. [45] The dynamics of maximum cell count in an individual generation for all 24 conditions have shown in FIG. S6, giving similar observations. P test for all 24 possible conditions has shown in FIG. S7.

### Clonal competition: survival clones and average size

Clonal competition has been observed in different types of stem cells in earlier experiments. [44, 45, 55, 56] It has observed that in clonal competition of mouse spermatogonial stem cells (activated with GFR*α*1 gene) the number of clones reduces with time, while average clone sizes increases with time. [45] Similar qualitative behavior of stem cell has been observed in other different experimental studies as well. [44, 55, 56] In our model, to study clonal competition, we have taken the simplest case such as non-competitive competition (see clonal competition section in material and methods). In this competition a stem cell stochastically divides and replaced by it’s daughter cells. We have started from 50 clones and observed that number of clones reduces with time, and dominated by fewer clones (FIG. 9A). These dominated clones (FIG. 9B) increase their sizes with time. FIG. 9C_1_-*C*_6_ show the number of clones at different-different events, and each colored slice represents the size of the clone. So the supports the hypothesis that in clonal competition number of clones reduces with time and average clone size increases.

**FIG 9:**
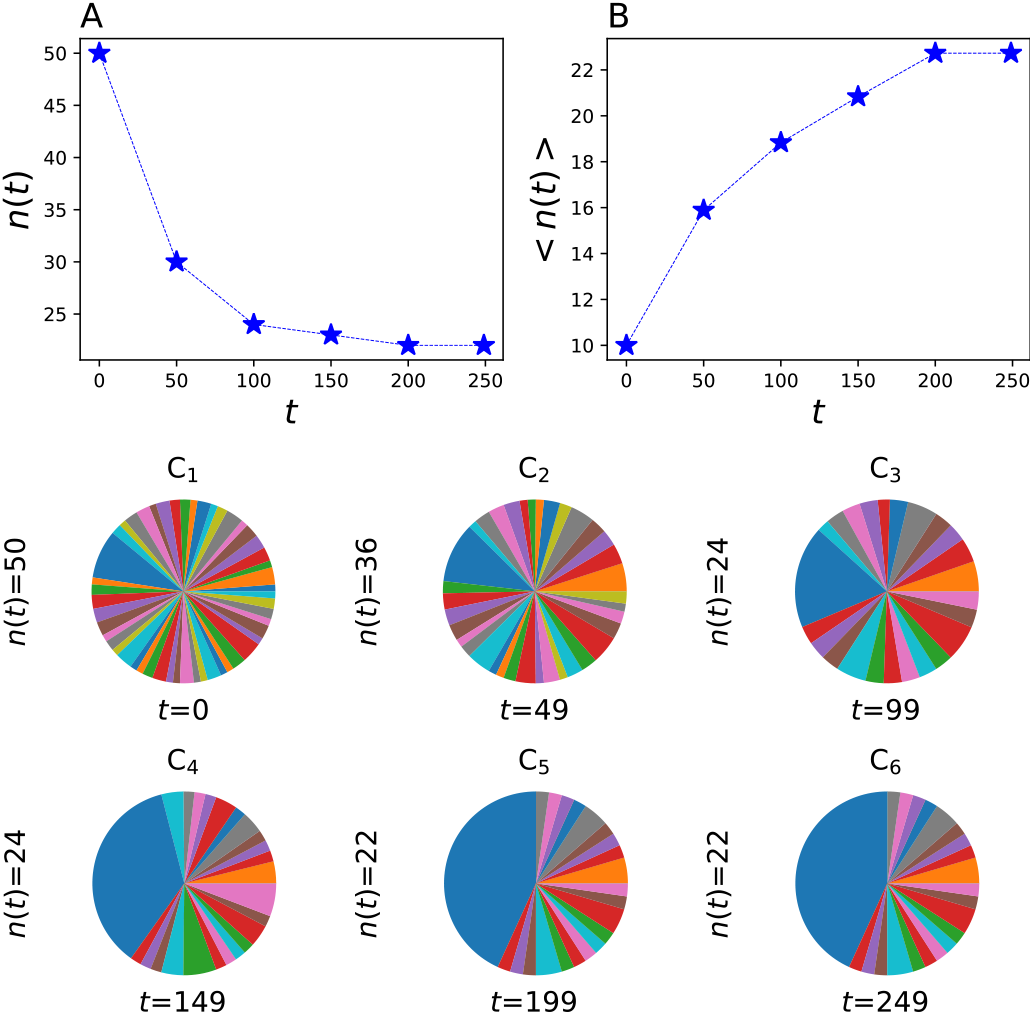
Clonal competition: A) displays decreasing behavior of clone numbers over event (time) showing that clonal competition is dominated by few clones. B) displays increase in average clone size over event (time). C_1_ -C_6_) pie chart represents the number clones at different events where each slice is showing the size of individual clone.

## Summary and discussion

A cell precisely balances between proliferation and differentiation to maintain homeostasis, and this diversity is regulated by two key regulatory mechanisms, intrinsic and niche. It is well known that stem cells are stochastically lost in tissue formation and replaced by daughter cells. However, it is not understood how a cell decides to either keep the same identity or adapt to a new identity. Key factors are suggested and verified by experiments that play a key role in cell fate decisions. Here we explore few key factors from a modeling perspective, which drives cell variability from a similar genetic level.

We purposed a population-based stochastic cell division model, starting from a single cell having three main components: nucleus position, segregation of transcription factors in daughter cells, and gene-regulatory network (GRN). Nucleus position decides plane of cleavage whether cell divides horizontally or vertically. After division, transcription factors are stochastically segregated in the daughter cells. Evolution of transcription factors is governed by GRN which has two transcription factors inhibiting each other and self-activating. GRN consists of three different attractors, each attractor corresponding to a particular cell state and differentiation is taken as transition from stem cell attractor to different attractor which is driven by internal/external factors. Here we have studied few questions such as the effect of factors leading to the stochastic nature of individual cell division, understanding short and non-terminating genealogies of stem cell, and universal clonal competition pattern such as dominance of few clones with increasing average size. The model leads to one key point that each cell at each generation makes a stochastic decision of cell division and differentiation. Our model results are in good agreement with experimental observations.

The model gives different types of genealogies with different range of terminal generation. Mainly two different types of genealogies, short and non-terminating (FIG. 4) are observed as in cell lineages experimental studies [15, 49–53]. Short genealogies has more symmetric trees (mainly in the first and second generation) and very less in non-terminating genealogies as increasing the tree size with different nodes reduces the chances of getting symmetric trees. Symmetric division leads to non-terminating whereas horizontal division gives short terminated genealogies. These observed genealogies suggest that probability distribution of cell might be a bi-modal distribution with two different modes. One mode corresponds to 0 stem cells, and second corresponds to non-zero stem cells. We have observed similar bi-modal probability distribution (FIG. 5A,B,C, FIG. 6, FIG. S1, and FIG. S2) of stem cells and differentiated cells, which is in very good agreement with Markov chain (FIG. 5G,H,I). We have observed that nucleus position towards basal enhances asymmetric division while towards apical leads to more symmetric division (FIG. 7) as observed in experimental studies [39]. This might be due to equal segragation of transcription factors (FIG. 7) in daughter cell as nucleus position moves towards apical. We have also observed that daughter cells having same volume and same population of transcription factors can lead to asymmetric division, and daughter cells having different volume and transcription factors can lead to symmetric division (because of intrinsic noise in evolution of GRN). Introducing noise in the nucleus position increases differentiated cells and reduces stem cell population (FIG. 6 and 7), which indicates that noise can alter the fate from symmetric division to asymmetric division. Noise also increases the number of generations and time duration in achieving a certain population by giving asymmetric division. This might explain how niche signalling maintain homeostasis through out the life by asymmetric division.

We have studied four different cases *horizontal, vertical, mixed*, and *random* cases with noise and no-noise in nucleus position. In the case of noise in the nucleus position, horizontal division leads to more differentiated cells in contrast, vertical division gives more stem cells. In the case of no-noise, both result in more stem cells, but vertical division has more stem cells than horizontal division (keeping *P* constant). In mixed case, stem cell increases and differentiated cell decrease as nucleus position shifts towards apical. Random case has no effect on cell population changing the nucleus position, while in case of noise, it produces more differentiated cells. These observations indicate that noise increases fraction of differentiated cell line and *mixed* case would be more suitable case to study further for cell-cell communication and clonal competition. Total cell number, contributed by each generation, shows an increased cell line followed by decreasing pattern after attaining a certain cell count. We also observed that cell dynamics in individual cell generation follow a Gaussian distribution with variable in maximum cell count (FIG. 8 and FIG. S4). The model simulation results are in good agreement with experimental studies [51, 54]. There are patterns which are similar in clonal competition of different types of stem cells. [44, 45, 55, 56] Such as decreasing pattern of number of clones, increasing pattern of average clone size, fraction of clones with *clone size/average clone size* at any instance of time, dominance of clonal competition by fewer clones, and others. [44] Here, we have also observed decreasing pattern of clone number (domination of fewer clones) and increasing pattern of average clone size (FIG 9) which are in consistence with hypothesis. Although, through out the model proportion of A, and B cell were observed similar which was due to having same parameter values in GRN. Different proportion of cell A and B are observed taking different values of *n*, and *m*, as same noise level will bias towards one differentiated cell attractor. Increasing initial volume of cell also leads to more stem cell and non-terminating cell linage tree.

The proposed stochastic model is capable of showing different dynamics of stem cell division, which are in good agreement with experiments. Self-renewal and differentiation both can be achieved by two different regulatory mechanisms intrinsic and niche. In this model, we did not include any external signal which could alter cell fate. It could be interesting to see how the external signal alters cell fate and how it affects cell proportion. External signal can be modeled by adding diffusive term in genetic regulatory network (GRN) [35–37]. It could be more interesting to see the clonal competition in different tissue as average clone size *<n*(*t*)*>* changes with tissue type such as 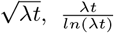, and *λt* in 1D, 2D, and 3D tissues respectively [44], *λ* being stem cell loss rate and *t* time.

## Author contributions

Conceptualization: AJ, SS, RR; Methodology: AJ, SS, RR; Analysis: AJ, SS, MZM, RR; Writing - original draft: AJ, SS, MZM, RR; Writing - review editing: RR; Supervision: RR.

## Supplementary information

Supplementary information available online.

## Appendix

### Rules of cell division

1. Gene regulatory network: The gene regulatory network consists two transcription factors *X*, and *Y* which inhibit each other and activate itself. The system exhibits three different attractors, each corresponds to a different cell type.

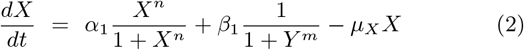

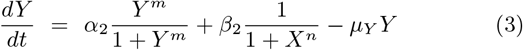

a). Type A (*X > Y*) b). Type B (*X < Y*) c). Type S (*X* ≈ *Y*) *X*, and *Y* are concentration of transcription factors, *α*_1_ = 0.6, and *α*_2_ = 0.6 are autocrine parameters, *β*_1_ = 0.3, and *β*_2_ = 0.3 are paracrine parameters, *n* = 3, and *m* = 3 are hill coefficient, *µ*_*X*_ = 0.3, and *µ*_*Y*_ = 0.3 are death rates of transcription factors *X*, and *Y* respectively.
2. Initiate with initial nucleus position *P*_0_ ∈ (0, 1), initial volume *V*_0_ of stem cell, and a tree with root S (Stem cell). We have taken *V*_0_ = 2 (initial volume of stem cell as root in each tree) through out the model.
3. Start with population of *X*_0_, and *Y*_0_ for regulatory network which corresponds to stem cell attractor (Step 1).
4. Simulate volume dependent Gillespie for *V* → 2*V*_0_ with *V* = *V*_0_*exp*^*λt*^, where 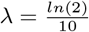.
5. Stochastic partition during cell division (Plane of cleavage and segregation of transcription factors):
  a. Pick the cell based on *P* (nucleus position) value.
  b. Generate random number r_1_ ∈ (0,1), if r_1_ *> P* : Horizontal division, else: Vertical division.
  c. Divide the cell into two daughter cells with volume *V*_1_, and V_2_ as volume reaches up to 2*V*_0_. If division is horizontal division then *V*_1_ will be proportional to *P* and *V*_2_ will be proportional to (1-*P*), else *V*_1_ = *V*_2_ = *V* /2 (vertical division).
  d. Generate the random number r_2_ ∈ (0,1), if 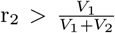 the one molecule of *X* will be going in cell volume *V*_2_ otherwise it will be going into cell volume *V*_1_. Repeat the process upto number of molecules of *X*. Same process for the population of molecules *Y*.
  e. Track the deterministic time of cell division (death time) of each daughter cell by 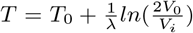. *T*_0_ is birth time of the cell, 2*V*_0_ is the final volume of the cell (which is always 4 in this model), and *V*_*i*_ is the initial volume of that cell.
6. Normalize the nucleus position of daughter cells after each division.

### Horizontal division case

a. If *P* 0 ≤.5 then nucleus position for first cell will be (say *P*_1_) 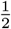, while for other cell it will be (say *P*_2_) 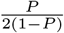.
b. If *P >* 0.5 then nucleus position for first cell will be (say *P*_1_) 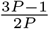, while for other cell it will be (say *P*_2_) 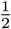.

### Vertical division case

a. No change in the nucleus position.

### Introducing noise in nucleus position

a. For cell 1: *P*_1_ = *P*_1_ + unifrom random number between - 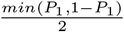to 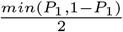.
b. For cell 2: *P*_2_ = *P*_2_ + unifrom random number between - 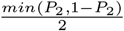 to 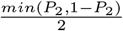. Note: Follow Step: 5b to check horizontal or vertical division case in each division.

7 Convert transcription factors population *X*, and *Y* into concentration by dividing volume *V*_1_, and *V*_2_ respectively. Update the tree by simulating ODEs for the cell type (to know which cell type it is).
8 After division track each cell [*T* +*τ* (next division time), T (birth time), *V, P, X, Y, parent node, daughter node, daughter fate*], and update genealogical tree.
9 Go to Step: 4 and repeat up to a desired generation of cell division.

