## Supplementary information for "A stochastic model of homeostasis: the roles of noise and nuclear positioning in deciding cell fate"

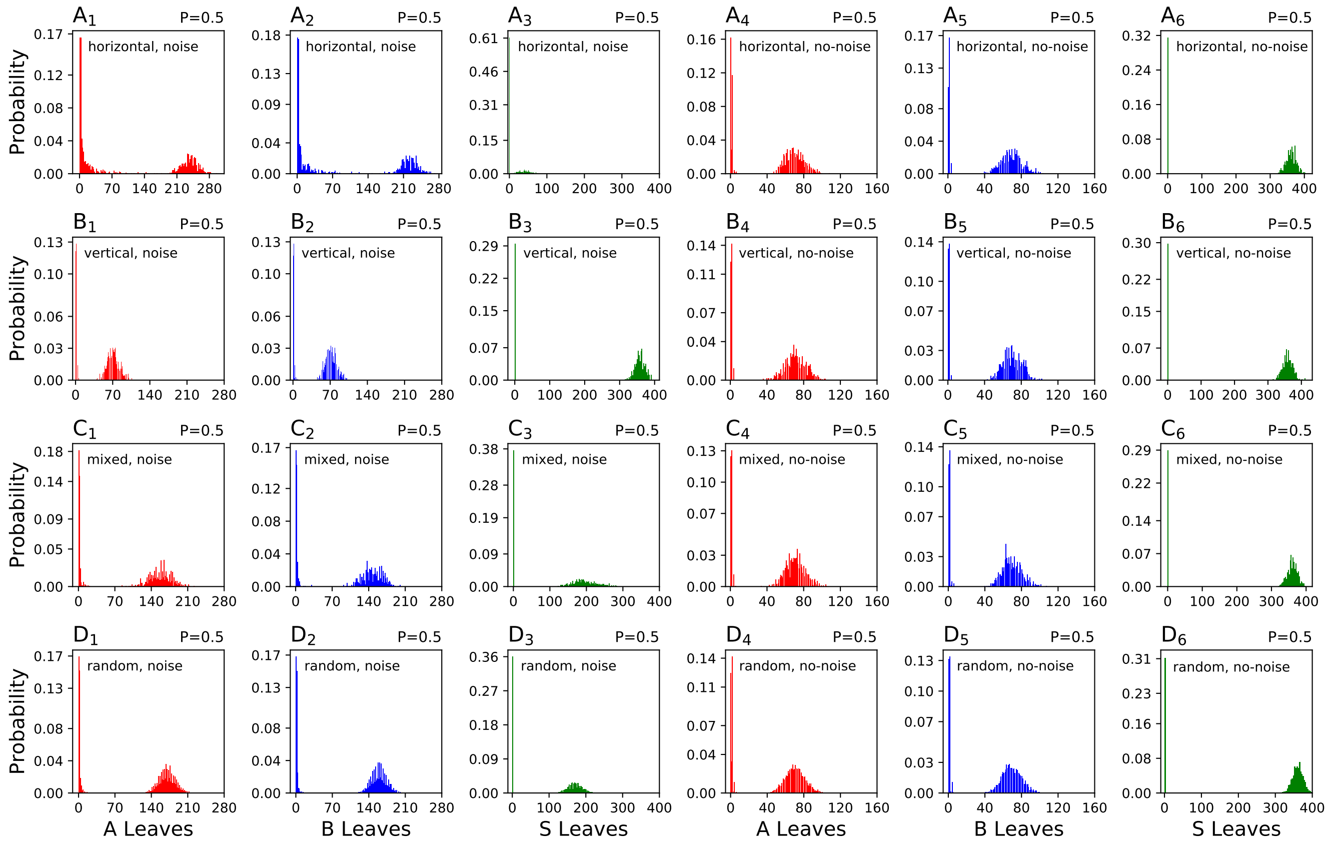


FIG. S1: **Probability distribution of cell types A, B, and stem cell S for nucleus position P=0.5**. Probability distribution of cell A (differentiated, red color), B (dierentiated, blue color), and S (stem cell as a leaf, green color) for all eight different cases asymmetry, symmetry, mixed, and random with noise and without noise in nucleus position, averaged over 5000 samples. P (right on the top of each panel) represents the nucleus position, and inside the panel, the first and second text represents the type of cleavage and noise in the nucleus position, respectively.

*Electronic address:

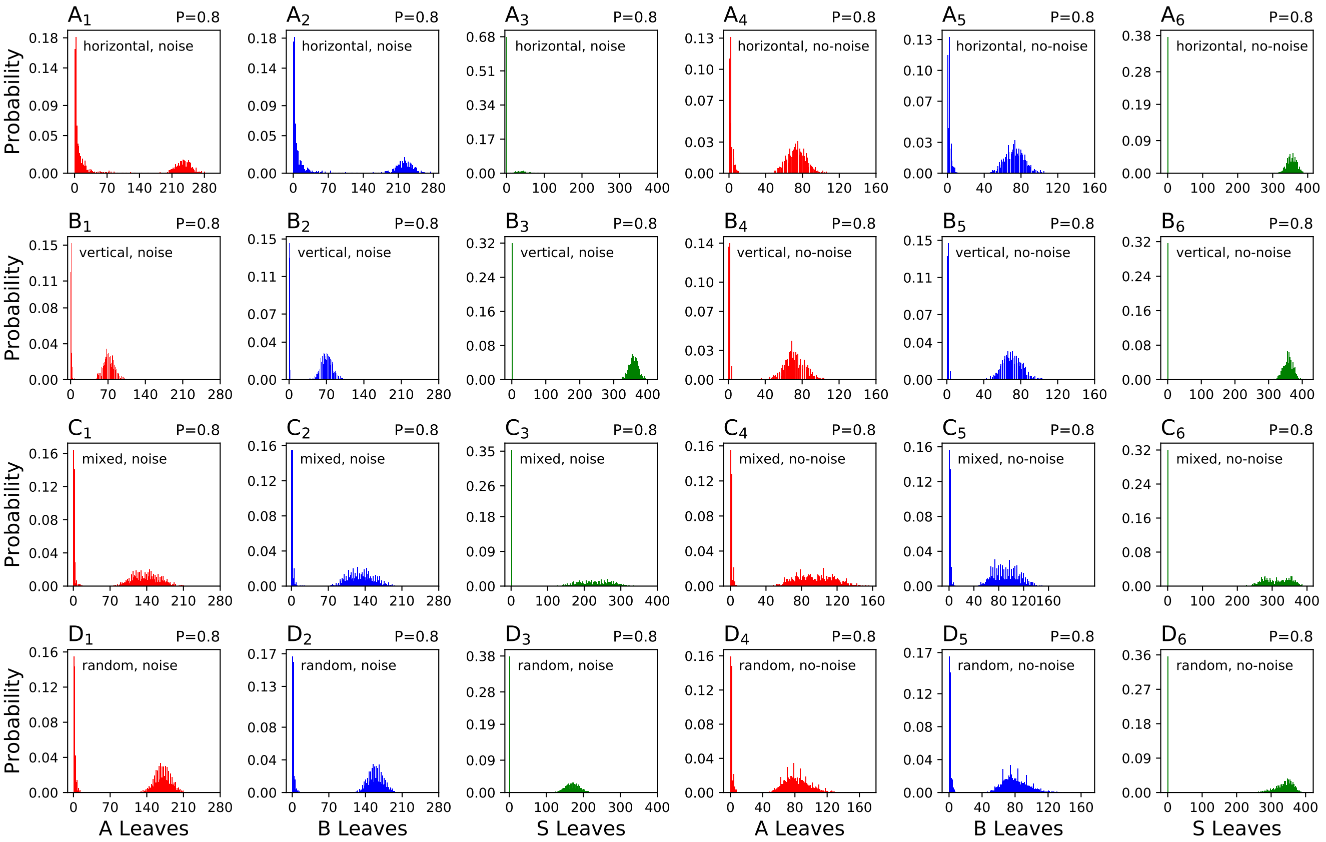


FIG. S2: **Probability distribution of cell types A, B, and stem cell S for nucleus position P=0.8**. Probability distribution of cell A (differentiated, red color), B (dierentiated, blue color), and S (stem cell as a leaf, green color) for all eight different cases asymmetry, symmetry, mixed, and random with noise and without noise in nucleus position, averaged over 5000 samples. P (right on the top of each panel) represents the nucleus position, and inside the panel, the first and second text represents the type of cleavage and noise in the nucleus position, respectively.

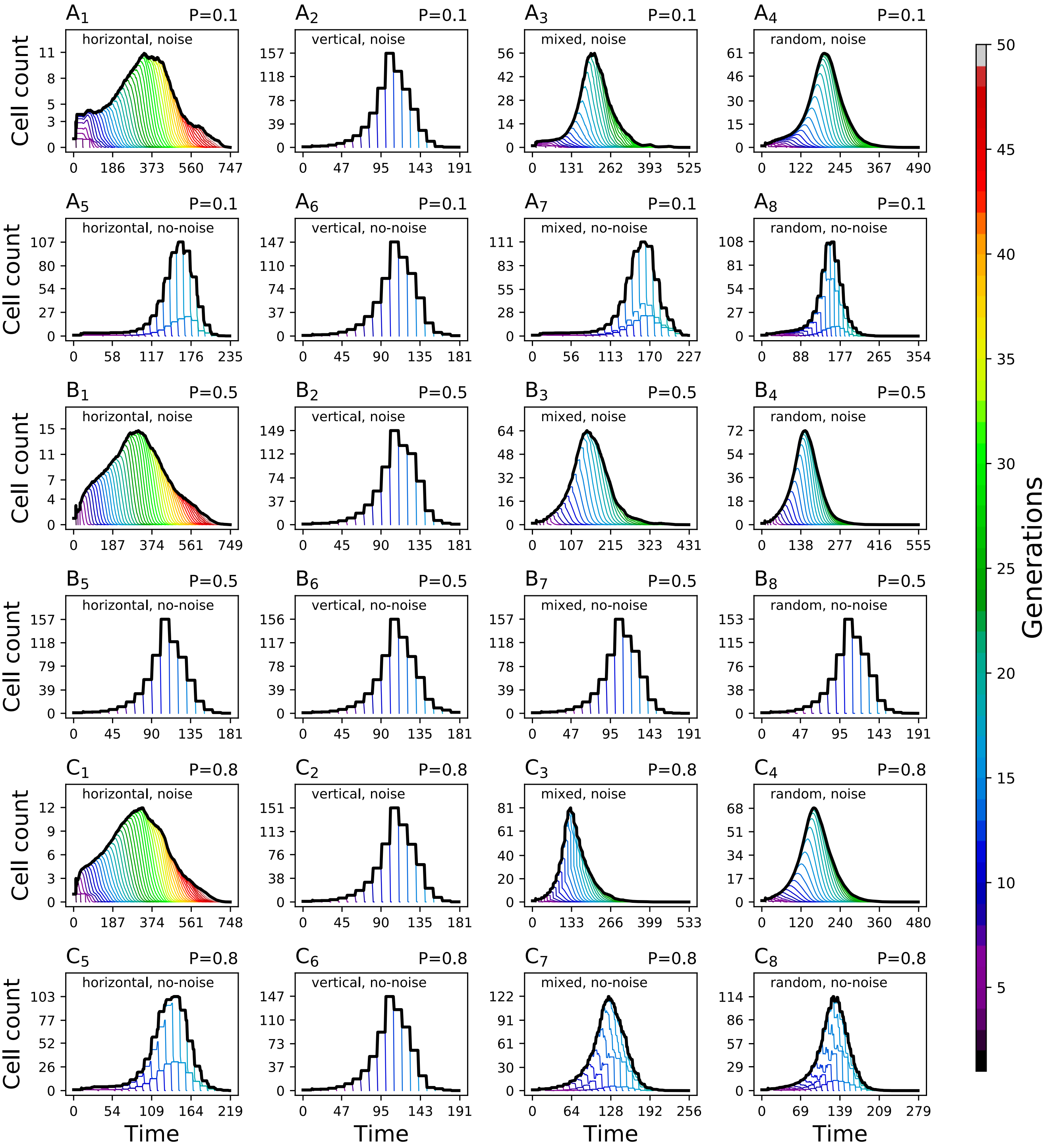


FIG. S3: **Population dynamics of cell**. Cell count in different generations (stacked plot) over time (averaged over 5000 samples). The thick black line represents the dynamics of total cell counts as a function of time, and color lines represent the dynamics of cell in different generations. The color spectrum represents color for each generation. P (right on the top of each panel) represents the nucleus position, and inside the panel, the first and second text represents the type of cleavage and noise in the nucleus position, respectively. In all the panels, we have plotted cell dynamics up to 50 generations.

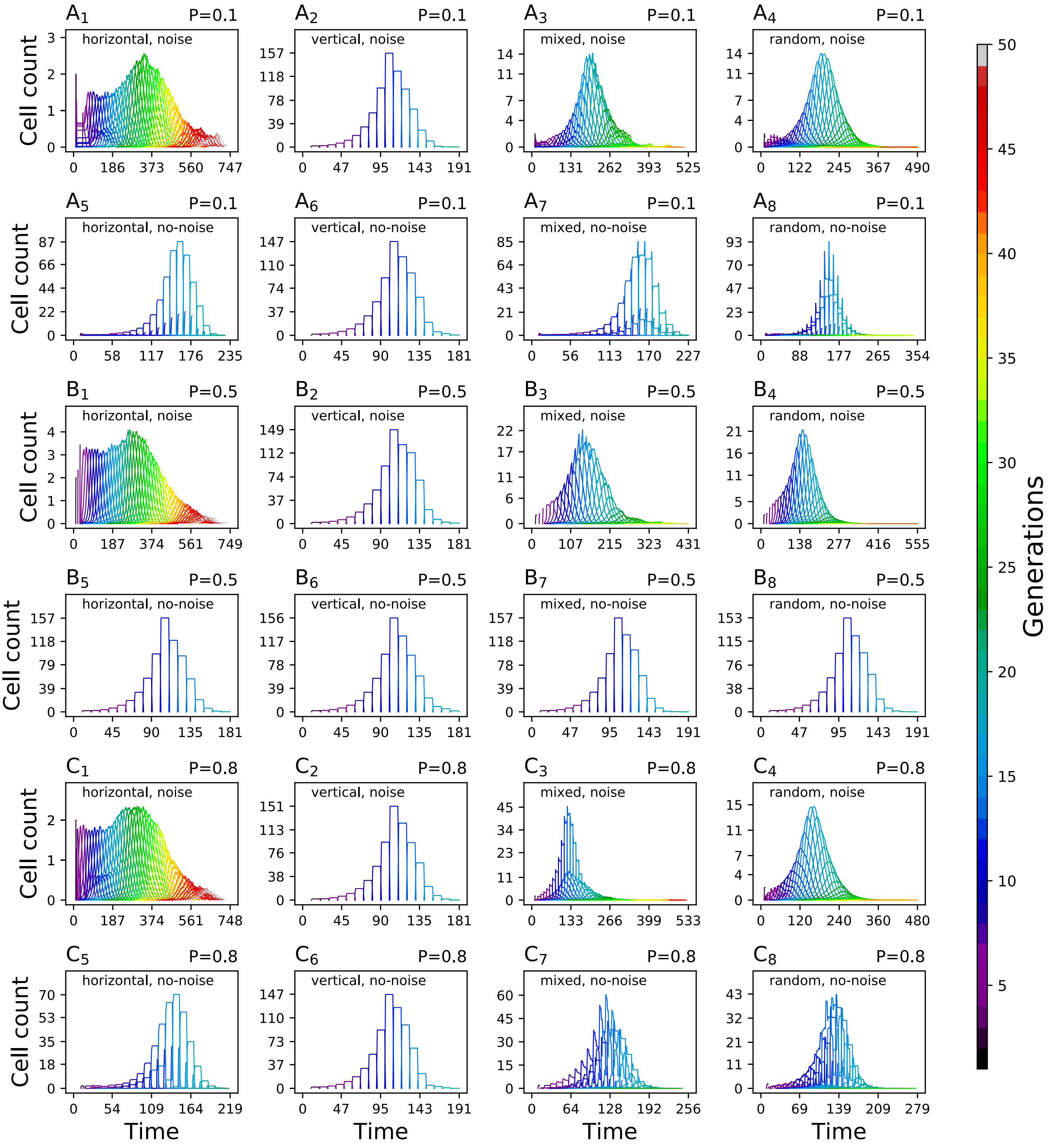


FIG. S4: **Population dynamics of cell in individual generation**. Cell dynamics in individual generation as a function of time which show Gaussian distribution. Color spectrum shows color for each generation. P (right on the top of each panel) represents the nucleus position, and inside the panel, the first and second text represents the type of cleavage and noise in the nucleus position, respectively. In all the panels, we have plotted cell dynamics up to 50 generations.

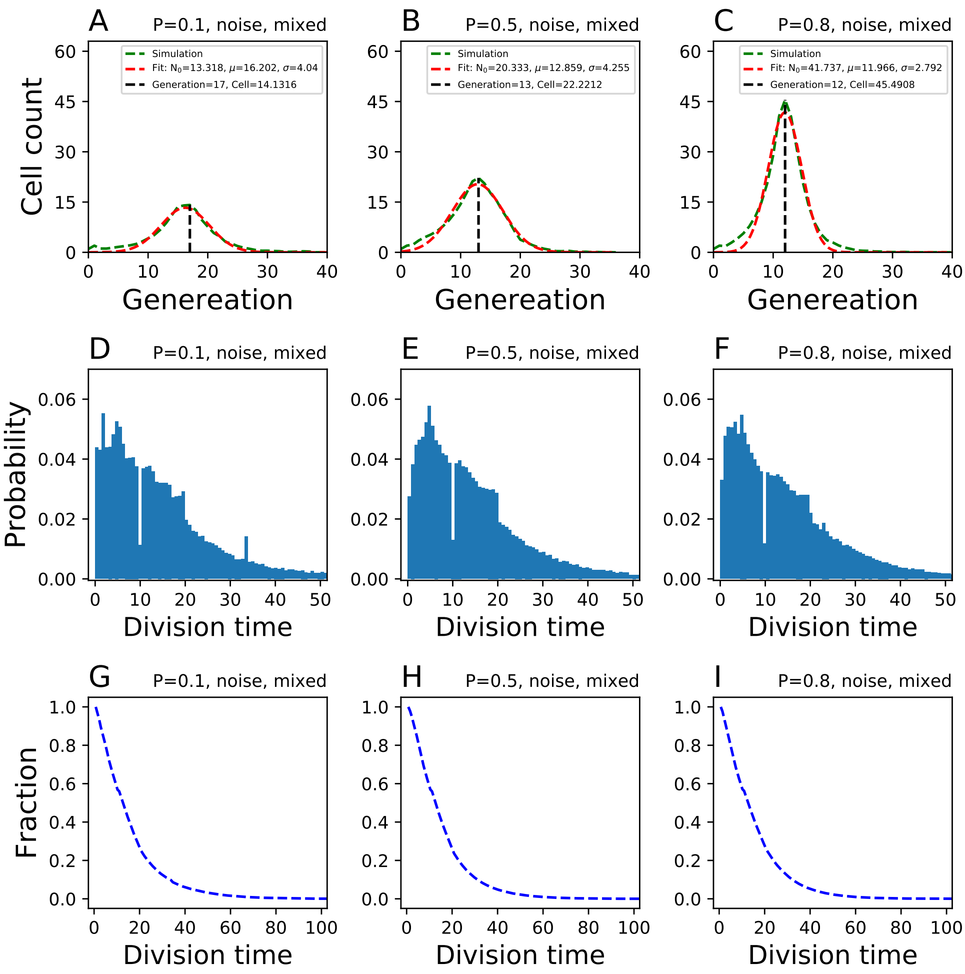


FIG. S5: **Decaying pattern of division time and division fraction**: In the top row, green dashed line represents the dynamics of maximum cell number in an individual cell generation, and red dashed line show Gaussian fit. Black vertical line corresponds to generation having maximum number of cells. Second row represents probability distribution of cells as a function of division time. Bottom row correspond to the division fraction of cell as a function of division time.

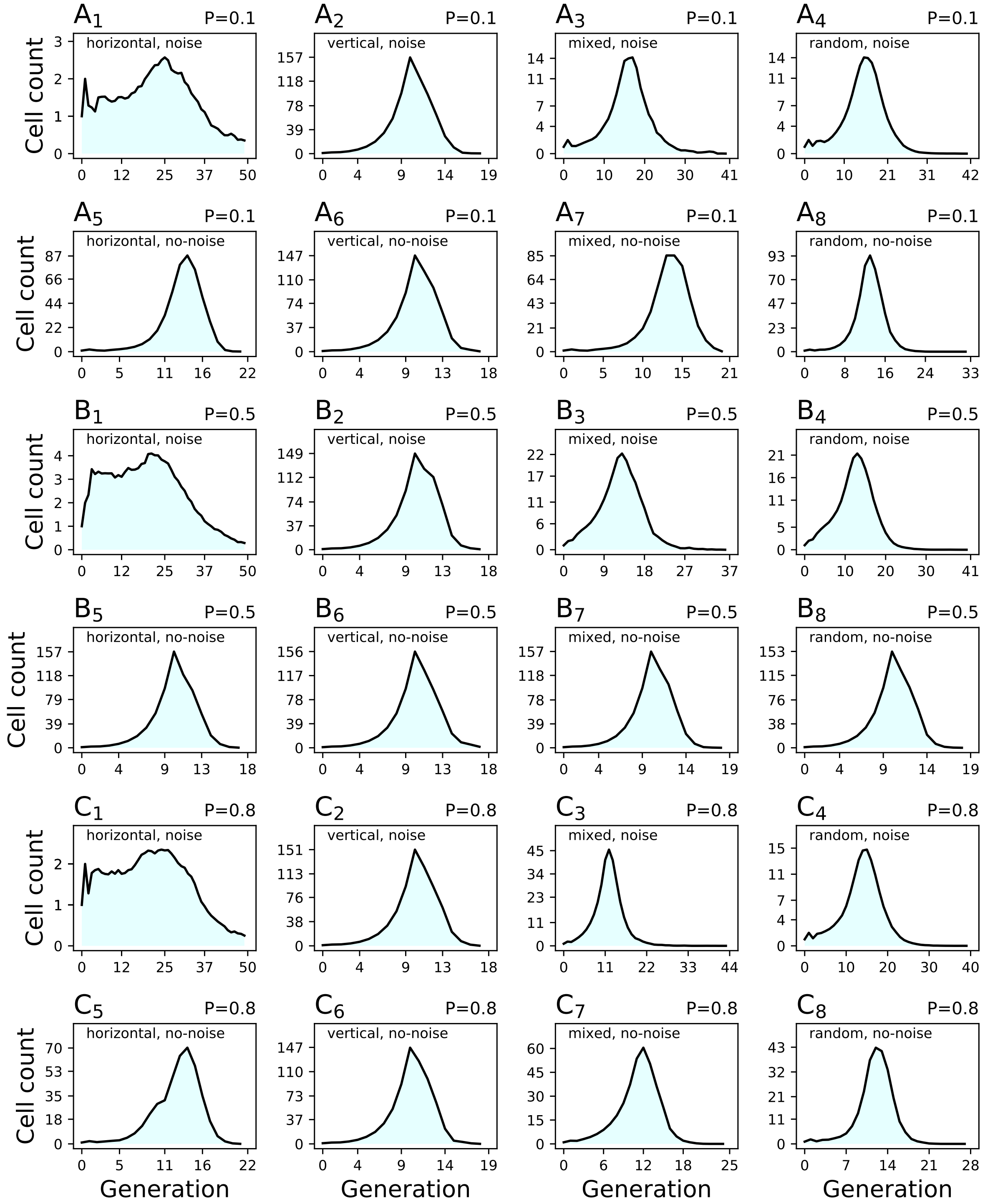


FIG. S6: **Population dynamics of cell over generation**. Distribution of maximum number of cells in individual generation over time (averaged over 5000 samples). P (right on the top of each panel) represents the nucleus position, and inside the panel, the first and second text represents the type of cleavage and noise in the nucleus position, respectively. In all the panels, we have plotted up to 50 generations.

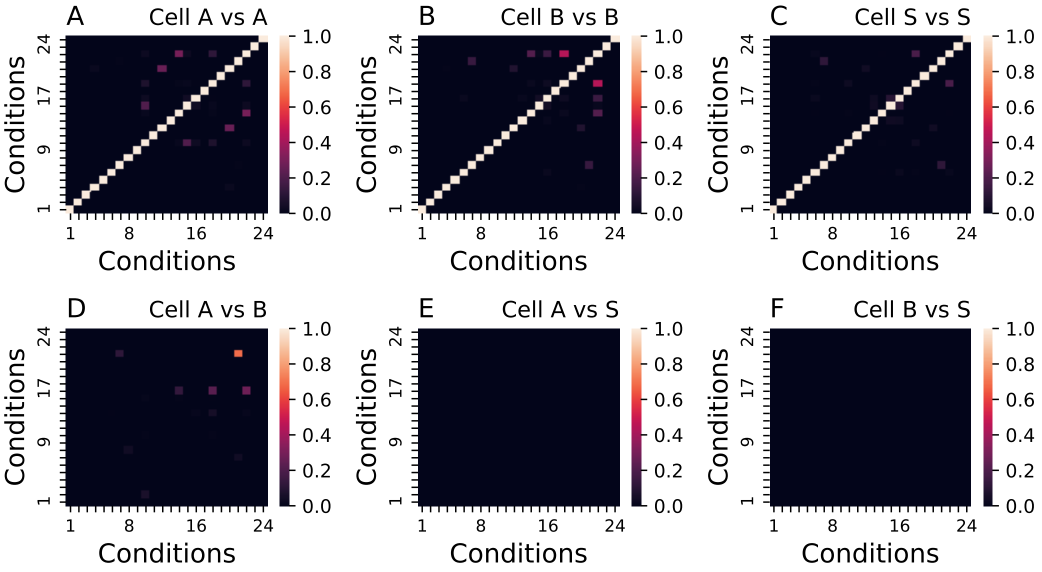


FIG. S7: **P-test**. P-test for possible 24 cases described in TABLE I. Each point on heat map represents P value corresponding to color bar.

TABLE I: Description of conditions used in FIG. S7

| **Condition** | **Description (nucleus position, division plane, noise)** |
|---|---|
| 1 | (0.1, horizontal, noise) |
| 2 | (0.1, vertical, noise) |
| 3 | (0.1, mixed, noise) |
| 4 | (0.1, random, noise) |
| 5 | (0.1, horizontal, no-noise) |
| 6 | (0.1, vertical, no-noise) |
| 7 | (0.1, mixed, no-noise) |
| 8 | (0.1, random, no-noise) |
| 9 | (0.5, horizontal, noise) |
| 10 | (0.5, vertical, noise) |
| 11 | (0.5, mixed, noise) |
| 12 | (0.5, random, noise) |
| 13 | (0.5, horizontal, no-noise) |
| 14 | (0.5, vertical, no-noise) |
| 15 | (0.5, mixed, no-noise) |
| 16 | (0.5, random, no-noise) |
| 17 | (0.8, horizontal, noise) |
| 18 | (0.8, vertical, noise) |
| 19 | (0.8, mixed, noise) |
| 20 | (0.8, random, noise) |
| 21 | (0.8, horizontal, no-noise) |
| 22 | (0.8, vertical, no-noise) |
| 23 | (0.8, mixed, no-noise) |
| 24 | (0.8, random, no-noise) |
